# Is *Aedes nr. albopictus* synonymous with *Aedes pseudalbopictus?*

**DOI:** 10.1101/2025.06.28.661682

**Authors:** Om P. Singh, Radhika Mittal, Rajnikant Dixit

## Abstract

A recent study reported a putative cryptic species of *Aedes albopictus*, provisionally named *Aedes nr. albopictus*, from Tripura, India, based on molecular data from mitochondrial cytochrome c oxidase subunit I (COI) and nuclear internal transcribed spacer 2 (ITS2) regions. However, our reanalysis, incorporating COI and ITS2 sequences of the co-occurring species *Aedes pseudalbopictus* and employing maximum likelihood phylogenetic analysis, albeit with a limited number of available sequences, suggests that this lineage is conspecific with *Ae. pseudalbopictus*. Given the morphological similarities and overlapping habitat use of *Ae. albopictus* and *Ae. pseudalbopictus* in India and Southeast Asia, we hypothesize that the reported cryptic species could potentially be attributed to *Ae. pseudalbopictus*. Our findings underscore the need for comprehensive taxonomic assessments, integrating morphological and molecular analyses of all sympatric *Stegomyia* species, to ensure accurate species delimitation and support effective vector surveillance and control.

## Background

Biswas *et al*. (2025) [1] recently identified *Aedes nr. albopictus*, a putative cryptic species sympatric with *Aedes albopictus* in Tripura, India, based on molecular analysis of mitochondrial cytochrome c oxidase subunit I (COI) and nuclear internal transcribed spacer 2 (ITS2) regions. Their findings underscore its possible role in arbovirus transmission, with significant implications for vector control strategies. Biswas *et al*. noted that *Ae. nr. albopictus* shares genetic similarities with cryptic *Ae. albopictus* lineages previously reported by Minard *et al*. (2017) in Vietnam [2] and Guo *et al*. (2018) in China [3], suggesting a broader regional presence of this lineage.

Biswas *et al*. asserted that *Ae. nr. albopictus* is “genetically distinct from *Ae. albopictus, Ae. subalbopictus, <u>*Ae. pseudalbopictus*</u>, Ae. flavopictus, Ae. aegypti*, and *Ae. vittatus*” [1] but their claim of genetic distinction from *Ae. pseudalbopictus* is unsubstantiated, as their phylogenetic analyses did not include genetic data *Ae. pseudalbopictus*. Biswas *et al*.’s assertion appears to be an assumption, as their study lacks both molecular sequences and detailed morphological comparisons with this species. This critical omission undermines the validity of their conclusion regarding *Ae. pseudalbopictus*, leaving open the possibility that *Ae. nr. albopictus* may be conspecific with or closely related to *Ae. pseudalbopictus*, particularly given their morphological and ecological similarities in shared habitats across India and Southeast Asia [4-6].

Recently, Hide *et al*. (2024) [7] also described *Aedes unalom* sp. nov. as a new *Stegomyia* species in Cambodia, distinguishing it from *Ae. albopictus* and other species including *Ae. pseudalbopictus* using COI, 5.8S-ITS2, and morphological traits, such as an interrupted white stripe on the hind femur [7]. While Hide *et al*. included *Ae. pseudalbopictus* sequences from their Cambodian collections, they did not include sequences from the cryptic lineages reported by Minard *et al*. and Guo *et al*. in their phylogenetic analyses of *Ae. unalom*, leaving the phylogenetic relationship between new *Ae. albopictus* lineage (designated as *Ae. nr. albopictus* by Biswas *et al*.) and *Ae. pseudalbopictus* unresolved.

Given the morphological and ecological overlap between *Ae. albopictus* and *Ae. pseudalbopictus*, we hypothesize that *Ae. nr. albopictus* and the cryptic lineages reported by Minard *et al*. and Guo *et al*. may be conspecific with *Ae. pseudalbopictus*. To test this, we conducted a maximum likelihood phylogenetic analysis, incorporating COI and ITS2 sequences of *Ae. nr. albopictus, Ae. pseudalbopictus* (five COI from BOLD Systems [8] originating from China [9] and Cambodia [7] and three sequences of ITS2 [7]), *Ae. unalom, Ae. albopictus*, and *Ae. aegypti* as an outgroup, to resolve their genetic relationships and clarify species boundaries.

## Methods

Sequences of COI and ITS2 for *Aedes* nr. *albopictus* from Tripura [1], *Ae. pseudalbopictus* from Cambodia [7] and China [9], *Ae. unalom* [7] were retrieved from GenBank. COI sequences of *Ae. albopictus*, and *Ae. aegypti* were obtained from VectorBase and ITS2 sequences from GenBank. To ensure data quality, flanking sequences potentially containing primer-derived region or sequencing artifact were trimmed in case of COI sequence with accession number JQ728197 [9]. Sequences for each marker were aligned independently using ClustalW in MEGA X v10.2.2 [10] with default parameters (gap opening penalty: 15, gap extension penalty: 6.66) and manually adjusted to ensure accuracy. Alignments were trimmed to uniform lengths for downstream analysis. Phylogenetic analysis was conducted using IQ-TREE2 [11] under a maximum likelihood framework. Nucleotide substitution models for COI and ITS2 datasets were selected with ModelFinder Plus based on the Akaike Information Criterion (AIC), Corrected AIC (AICc), and Bayesian Information Criterion (BIC). For the COI dataset, the AIC and AICc favored TVM+F+G4, but the K3Pu+F+G4 model was selected based on BIC, incorporating unequal base frequencies (+F) and gamma-distributed rate variation with four categories (+G4). For the ITS2 dataset, the AIC favoured K3P+I, while AICc and BIC selected K2P+G4, which was chosen as the best-fit model per BIC, incorporating Kimura’s two-parameter model with gamma-distributed rate variation (+G4).

## Results and Discussion

Our phylogenetic analysis revealed that *Aedes nr. albopictus*, along with cryptic lineages reported by Minard *et al*. and Guo *et al*. cluster with *Aedes pseudalbopictus* for both loci—COI and ITS2 loci (Figure 1, panel A and B, respectively). Furthermore, the ITS2 sequences of *Ae. nr. albopictus*, despite the known intragenomic variation in ITS2, were 100% identical to those of *Ae. pseudalbopictus* reported from Cambodia [7]. Notably, both studies [1,7] identified two identical haplotypes each differing only in a number of GC repeats (microsatellite), a pattern also observed in ITS2 of other mosquito species, *Anopheles stephensi* [12]. These findings strongly suggest that the proposed cryptic species may be *Ae. pseudalbopictus*. Further, our inclusion of *Ae. unalom* sequences in analysis confirms a distinct clade from *Ae. albopictus* and *Ae. pseudalbopictus*/*Ae. nr. albopictus* (Figures 1), supporting its status as a novel species.

**Figure 1.**
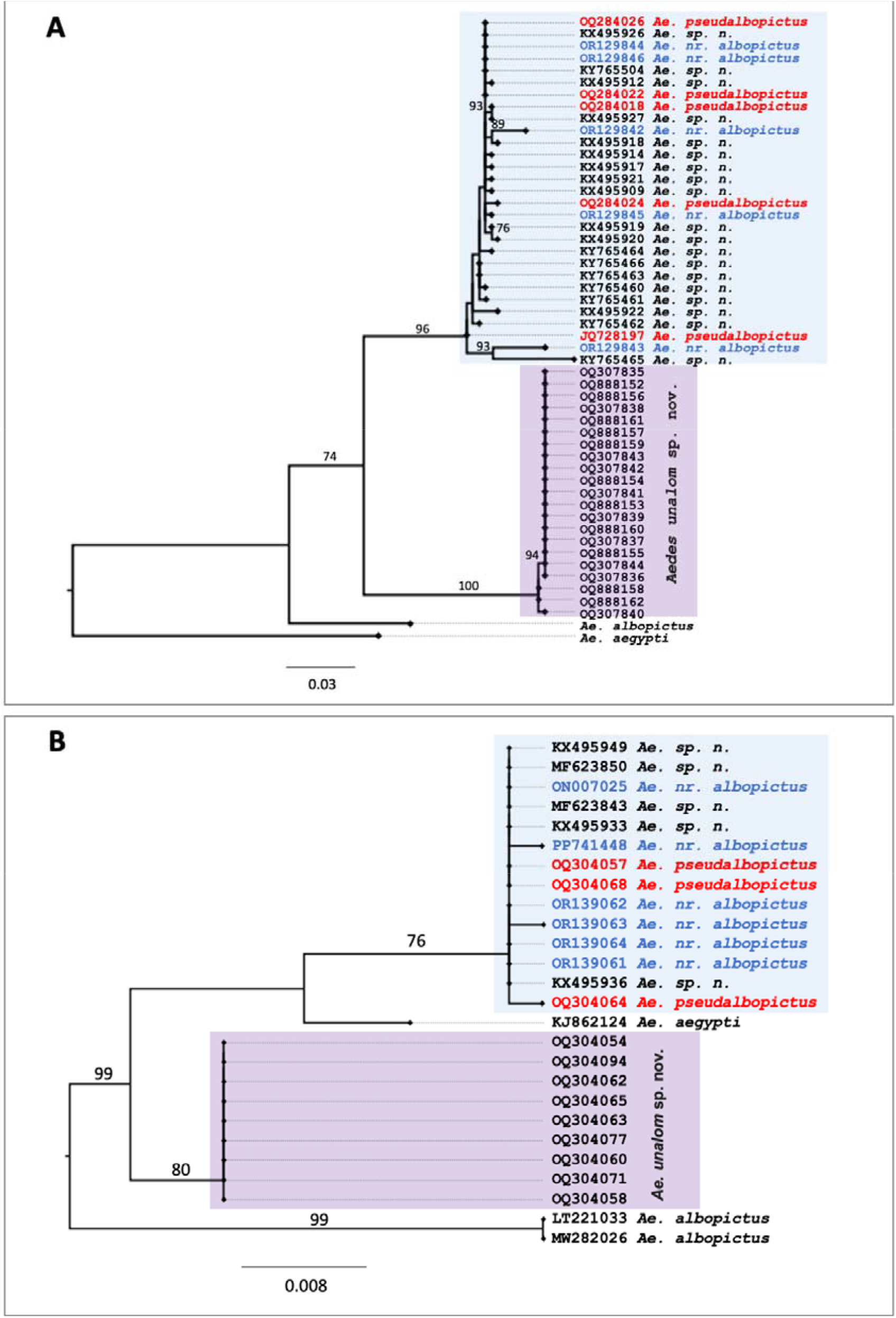
Maximum likelihood phylogenetic trees based on COI (panel A) and ITS2 (panel B) sequences of Aedes species. Ae. nr. albopictus from Tripura clustered with Ae. pseudalbopictus from Cambodia and China in both trees. Only bootstrap values ≥70% (1,000 replicates) are shown for clarity; scale bars indicate substitutions per site.

The misidentification of *Ae. pseudalbopictus* as *Ae. albopictus* likely stems from their morphological similarity. Huang (1968) [5] described subtle diagnostic characters, such as the presence of broad, flat white scales on the lateral margin of the scutum in *Ae. albopictus*, contrasted with narrow, curved white scales in *Ae. pseudalbopictus*. These differences are easily overlooked without meticulous examination. Additionally, Huang [5] noted that *Ae. pseudalbopictus* was frequently mistaken for *Ae. albopictus*, a confusion compounded by their sympatric distribution [5-6]. While Hide *et al*. explicitly identified *Ae. pseudalbopictus* in their collections using standard morphological taxonomic keys [14-15], the studies reporting new species [1-3] lacked specific details on the identification keys used to identify mosquito species. This lack of detailed methodology in those studies, combined with our phylogenetic findings, suggests that *Ae. pseudalbopictus* specimens may have been previously classified as a novel cryptic species of *Ae. albopictus*. On the other hand it may be possible that Wang *et al*. (2012) or Hide *et al*. (2024) misidentified their specimens as *Ae. pseudalbopictus* but their consistent morphological and molecular differentiation of *Ae. pseudalbopictus* makes this less likely.

The misidentification of *Ae. pseudalbopictus* as a cryptic species of *Ae. albopictus* has significant implications for vector surveillance and control in dengue- and chikungunya-endemic regions. *Ae. albopictus* is a well-documented vector, while the vectorial capacity of *Ae. pseudalbopictus* remains less studied. Misidentification could lead to inaccurate assessments of vector distribution, abundance, and behaviour, potentially skewing epidemiological models and control strategies. For example, differences in insecticide resistance or host-feeding preferences between *Ae. albopictus* and *Ae. pseudalbopictus* could affect the efficacy of interventions. Our analysis, by robustly confirming the conspecificity of *Ae. nr. albopictus* with *Ae. pseudalbopictus* and clarifying *Ae. unalom*’s distinctiveness, underscores the need for integrated taxonomic approaches that combine multi-locus molecular data with rigorous morphological examinations based on established keys, such as those by Huang (1972) [4] and Barraud (1934) [6].

While our analysis is robust, it is limited by the small number of *Ae. pseudalbopictus* sequences available (five COI and three ITS2 sequences); however, the consistent clustering of *Ae. nr. albopictus* with *Ae. pseudalbopictus* across both loci provides strong support for our conclusions. Additional sequencing of *Ae. pseudalbopictus* populations from India and Southeast Asia would strengthen our conclusions. To prevent taxonomic errors, we propose a standardized framework for *Stegomyia* species delimitation: multi-locus sequencing including mitochondrial and nuclear markers; morphological examinations using established keys; comprehensive sampling of all sympatric *Stegomyia* species; and, integration of ecological data to understand species boundaries. Special attention is required when analyzing rDNA, particularly ITS2, due to intragenomic variation due to indel or SNPs [12, 16-17]. Such variation can lead to ambiguous sequence chromatograms or alignment errors, potentially inflating perceived genetic divergence. Hide *et al*. have also reported mixed bases in ITS2 of *Ae. pseudalbopictus* which can be explained by presence of indel caused by variation in number of GC-repeats. This framework would enhance taxonomic accuracy and support effective vector control in regions where *Stegomyia* species are public health threats.

## Conclusion

The evidence presented here indicates that the reported new cryptic species *Aedes nr. albopictus* is conspecific to *Ae. pseudalbopictus*. This finding highlights the critical need for rigorous taxonomic studies that integrate morphological examinations with comprehensive molecular analyses, including all known sympatric species, before proposing new species. Accurate species identification is essential for effective vector surveillance and control, particularly for *Ae. albopictus* and *Ae. pseudalbopictus*, which are vectors of dengue and chikungunya in endemic regions.

## Author contributions

OPS: data analysis and writing first draft; RM: data analysis; RD: contributed to the manuscript.

## Competing Interests

The authors declare that they have no known competing interests

## Acknowledgement

The authors acknowledge the supportive research environment and resources provided by the Indian Council of Medical Research (ICMR) and the National Institute of Malaria Research (NIMR) that contributed to the development of the ideas presented in this manuscript. The views expressed are those of the authors and do not represent the official stance of ICMR or NIMR.

## Notes

### Competing Interest Statement

The authors have declared no competing interest.

